# A memristor-based neuromodulation device for real-time monitoring and adaptive control of *in vitro* neuronal populations

**DOI:** 10.1101/2021.05.20.444941

**Authors:** Catarina Dias, Domingos Castro, Miguel Aroso, João Ventura, Paulo Aguiar

## Abstract

Neurons are specialized cells in information transmission and information processing. Following this, many neurologic disorders are directly linked not to cellular viability/homeostasis issues but rather to specific anomalies in electrical activity dynamics. Acknowledging this fact, therapeutic strategies based on direct modulation of neuronal electrical activity have been producing remarkable results, with successful examples ranging from cochlear implants to deep brain stimulation. Development on these implantable devices are hindered, however, by important challenges: power requirements, size factor, signal transduction, and adaptability/computational capabilities. Memristors, nanoscale electronic components able to emulate natural synapses, provide unique properties to address these constraints and their use in neuroprosthetic devices is being actively explored. Here we demonstrate for the first time the use of memristive devices in a clinically relevant setting where communication between two neuronal populations is conditioned to specific activity patterns in the source population. In our approach, the memristor device performs a simple pattern detection computation and acts as a synapstor capable of reversible short-term plasticity. Using *in vitro* hippocampal neuronal cultures, we show real-time adaptive control with a high degree of reproducibility using our monitor-compute-actuate paradigm. We envision very similar systems being used for automatic detection and suppression of seizures in epileptic patients.

## 1. Introduction

Neurological disorders are a major cause of death and disability worldwide, and their burden in society continues to increase with population ageing and growth.^1^ Today’s therapeutic strategies still rely heavily on pharmacological approaches, with important problems regarding non-specificity and side effects. Furthermore, progress has been notably slow in discovering new drugs for diseases such as Parkinson’s, Alzheimer’s or epilepsy. Recognizing that neuronal function is intimately related with electrophysiology, attention is steadily growing towards a different therapeutic strategy: direct modulation of neuronal electrical activity. Deep brain stimulation for Parkinson’s disease, spinal cord stimulation for chronic pain, and cochlear implants for hearing loss are examples of success stories demonstrating the potential of this approach.

Neurotechnologies for therapies based on electrical modulation have, however, important challenges that still need to be addressed. These can be summarized in four keywords: size, power, interface, and computational capabilities. Two fundamental constraints in implantable medical devices are, naturally, their size and the power signature. In particular, the cost, burden, risk (e.g., infection) and idiosyncrasy of surgeries for battery replacement cannot be underestimated. The neurons-electronics interface (both signal and physical) is also a crucial challenge. Neurons and conventional electronics do not “talk” in the same electrical amplitudes and frequencies, limiting the effective (and direct) communication between the two systems. Noteworthy, there is also a scale mismatch between the neuronal structures (cell bodies and/or nerves) and the devices’ electronics. A last but also fundamental challenge has to do with the need for adaptive computation capabilities in the stimulation devices to cope with the dynamic nature of neuronal activity. A device that is unable to dynamically respond to changes in neuronal activity patterns falls short in its therapeutic potential and effectiveness.

It is only natural to address these neuroprosthesis challenges by mimicking key properties of the nervous system’s solution to the interface between neurons: the synapse. In this respect, memristive devices^2–7^ are a promising candidate to play the role of artificial synapses and integrate, as a core component, neuroprosthesis systems. Memristors are electrical components whose present conduction state depends on the voltage that has been previously applied to them. As (sub-)nanoscale two-terminal devices, their small feature size (<100 nm) enables high-density integration architectures and hardware implementation of powerful signal processors such as artificial neuronal networks. Memristors properties rely on the combination of materials used, namely the creation and rupture of nanoscale conductive filaments (e.g., Ag, Cu) under applied bias in metal-insulator-metal structures.^8–12^ Besides the low operation power involved in these mechanisms, the conductance state is also non-volatile staying unchanged when the power supply is removed, with additional benefits regarding power consumption. Among the most studied materials such as metal oxides (e.g., TiO_2_, HfO) and semiconductors (e.g, Si), the chalcogenides family (e.g., GeSe, Ag_2_S) have shown promising neuromorphic properties.^13–15^

The dynamics of a memristor’s response to stimulation is analogous to that of synapses, presenting learned transitions from low to high conductivity states, and vice-versa (**Figure 1A**). Importantly, the timescale of these transitions can be made analogous to the plasticity timescale of learning mechanisms in the brain, such as synaptic short-term plasticity (STP) which act on the range from seconds to minutes. Furthermore, memristors can be integrated in the back-end-of-line of complementary metal-oxide semiconductor (CMOS) technologies, in parallel with neuronal probe manufacturing.^16^ This means that memristive technology can leverage on the recent developments on high-density microelectrode arrays (MEAs) and neuronal probes carrying hundreds to thousands of recording/stimulating electrodes at an unprecedented spatiotemporal resolution.^17^

**Figure 1.**
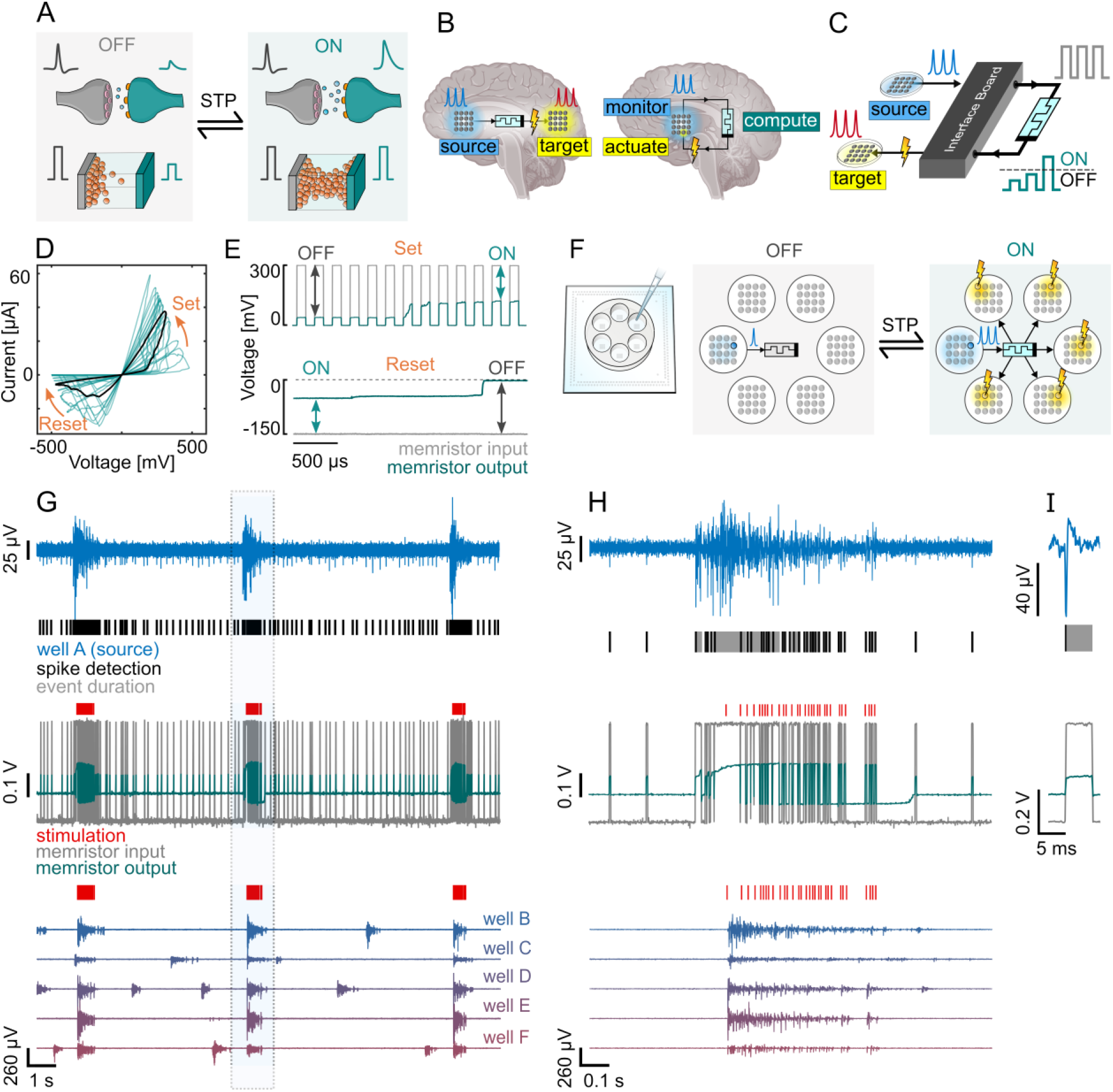
Real-time monitoring and adaptive control of neuronal populations using memristors. **A** Biological and memristive synapses share analogous short-term plasticity (STP), dynamics interchanging between low (OFF, left) and high (ON, right) conductance states as a function of the activity history. **B** Envisioned neuroprosthetics *in vivo*. The source and target neuronal population can be distinct (left) or be the same (right) providing either way an effective monitor-compute-actuate paradigm through the memristors-based device. **C** *in vitro* implementation used in this work, with a hardware interface bridging neuronal and memristor signals. **C** Memristor hysteretic loop showing “set” under positive voltage and “reset” under negative phases [*V*(*t*) = 0.5 sin(100*t*)]. The multiple cycles show the inherent variability of these type of devices. **E** Short-term plasticity dynamics with potentiation for repetitive positive pulsed (300 mV, 100 μs) stimuli (top) and recovery under constant negative (−150 mV) stimulus (bottom). **F** Schematic representation of the *in vitro* setting, integrating 6 independent neuronal cultures in a 6-well MEA (left) and a memristor to monitor the source population in real-time and dynamically modulate the activity of five target populations (right). **G** Representative example of the real-time neuronal monitoring and modulation performed by the *in vitro* memristive interface. Actuation upon the target population is gated by sustained/consistent high-level activity at the source electrode (blue, top). The detected spikes (black traces, top) are converted to pulses and fed to the memristor (grey, middle). The memristor output (green, middle) increases when the source bursts, changing to the ON state. Consequently, this triggers electrical stimulation (red traces, middle and bottom) at the target population, which is therefore modulated to burst with the source. The recordings shown for the targets (bottom) are from electrodes neighbouring the stimulation electrode. When the source bursts end, the memristor transitions back to the OFF state, and stops propagating the source spikes to the target. **H** Zoom of panel F evidencing the OFF-ON transition at the beginning of the source burst and the ON-OFF transition at the end. **I** Detail of a source neuronal spike and the associated detection performed by the interface board in real-time hardware (top). The spike creates a pulse that is applied to the memristor (bottom).

The recognition of the exceptional combination of properties in memristors have fostered important proof-of-concept studies demonstrating the feasibility of direct neuron-memristor connection.^18,19^ Memristors not only can be made to respond to neuronal activity, they can also act as an effective interface between biological and artificial neurons, implemented either by software or using very large-scale integration hardware.^20^ A unidirectional, activity dependent, direct coupling between two neurons in brain slices has also been recently achieved via organic memristive devices.^21^ In this study, however, the memristive devices only progress from a low to high conductance state (linking the neurons), limiting the functional usefulness of the coupling system. Neuronal activity in these systems is recorded/stimulated using either MEAs^18,19^, patch-clamp pipettes^21^ or a combination of the two.^20^ Memristors not only emulate fundamental synaptic properties, they can also be combined to achieve non-trivial computations. Important examples, with particular interest for memristors-based neuroprosthesis, include the use memristors to perform real-time processing of neuronal spikes^16^, or the use of memristor arrays to implement the filtering and identification of epilepsy-related neural signals.^22^

Although these proofs-of-concept studies have been covering important separate aspects of the potential role of memristors in implantable neuroprosthetic devices, to the best of our knowledge, no work has yet demonstrated the use of memristors in a fully functional configuration and in a clinically relevant setting. In this work, we take as motivation the development of closed-loop neurostimulators for intractable epilepsy. We believe such devices should perform three core tasks in a monitor-compute-actuate paradigm (**Figure 1B**): 1) online monitoring of a neuronal population prone to seizures, 2) real-time seizure detection, and 3) stimulation of specific (inhibitory or interfering) neuronal populations to suppress seizure progression. It has already been well-established that electrical stimulation can ameliorate epilepsy.^23^ To produce our control setting, we developed a memristor-based interface (fully implemented in hardware) capable of performing real-time monitoring and adaptive coupling between two neuronal populations (**Figure 1C**). Our system establishes direct communication between neuronal populations (not individual neurons) and *in vitro* experiments are carried out with neurons from the hippocampus, a brain region frequently involved in epilepsy. The reversible short-term memory dynamics of the memristors are used both for the detection of networks bursts (synchronous high-level activity in the neuronal population, typical in epileptic seizures) and for the gating of electrical stimulation to the target neuronal population.

## 2. Results and Discussion

In this work we used commercial memristors composed of stacks of W/C+Ge_2_Se_2_/SnSe/Ge_2_Se_3_ Mix/Ag/Ge_2_Se_2_ Adhesion/W.^24^ These memristors rely on the movement of silver ions (Ag^+^) into channels within the active layer, which has been doped with carbon to enhance and optimize their properties, and are characterized by low power binary switching. During the initial step of electroforming, under an applied positive voltage on the top electrode, Sn ions from the SnSe layer are generated and diffused into the active Ge2Se3 layer, where a metal-catalyzed reaction distorts the glass network to provide conductive channels for the movement of Ag^+^. Since the amount of Ag within the channel determines the resistance of the device, the resistance is then tunable by the movement of silver into or away from these channels by applying positive or negative potential, respectively (**Figure 1D**).^25^ Notably, these type of memristors are capable of short-term memory dynamics. Of particular importance for this work is the reversibility of the ON-OFF conductance transitions. With conditioning positive pulses and constant negative voltage, we were able to produce interchangeable “set” and “reset” transitions, respectively, with adequate timescales for a neurobiology setting (**Figure 1E**). Aiming at population’s level control, our designed system relies on MEAs to record and modulate neuronal activity, thus allowing long-lasting viable interfaces (as opposed to patch-clamp electrodes). As to discriminate the memristive device’s specific contribution to neuronal activity modulation, we used neuronal cultures on MEAs with a 6-well configuration, forcing the existence of 6 independent neuronal populations (**Figure 1F**). Communication between the source population/well and the remaining populations/wells was mediated by the memristor, which selectivity commanded electrical stimulation on the target populations according to the patterns of activity detected in the source population.

Currently there are still no memristors available working on a voltage/current range compatible with a direct connection to neurons (μV and nA). As such, and for now, instead of a direct/passive circuit neurons-microelectrode-memristor-microelectrode-neurons, our system includes hardware (e.g. amplifiers, electrical stimulator) to translate between microelectrodes and memristors’ voltage amplitudes (**Figure S1**).

### 2.1 Memristive interface detects and selectively responds to network bursting activity in real-time

Initially, the memristor is in a non-conductive state (OFF), analogous to a weak synaptic connection. Bursting activity at the source electrode induced a gradual increase in the memristor conductivity, changing its state to ON. When the memristor was ON, source spikes triggered electrical stimulation in the target population, inducing them to fire with the source population (**Figure 1G,H**). Importantly, the stablished neuronal coupling/modulation is reversible and the connection is dissolved when the spiking rate of the source decreases back to baseline activity. Note that the evolution of the memristor’s state is automatic and unsupervised, which means that no additional system is used to actively change its conductance - the memristive interface does the network burst detection computation autonomously.

The memristor receives a pulse for every spike in the source electrode (**Figure 1I**). A conditioning circuit (Experimental Section and **Figure S1**) was fine-tuned prior to the experiments to guarantee that the different patterns of activity exhibited by the neurons (patterns of received pulses) induce the memristor to interchange between its conductive states, ensuring the desired selective response to bursts. We lowered the amplitude the arriving signal to assure that each pulse had a positive amplitude, lower than the memristor’s “set” threshold. A pulse with an amplitude above the “set” threshold would immediately transition the memristor state to ON with a single source spike, eliminating any computation capabilities. When no spikes were being sent, the voltage across the memristor was negative with a small amplitude, slowly changing the resistance back to OFF. These parameters - amplitude of the pulse and negative baseline level - were optimized *a priori* and kept unchanged for all experiments. This was performed taking into consideration the natural stochasticity of the memristive behaviour. In terms of material dynamics, the positive pulses are gradually and cumulatively creating a metallic filament across the device, whereas the negative constant voltage is slowly dissolving the filament. Note that the latter would not be needed in the case of volatile memristors.

### 2.2 Memristive interface promotes robust coupling and modulation of neuronal populations

The memristor communicates with the neuronal cultures through a source electrode in the source well, and a stimulation electrode in each target well (**Figure 2A**). The bursting periods detected by the memristor are associated with network wide events that dominate the cultures activity (**Figure 2B**), referred as network bursts (NB). Although the memristor responds to the spikes from a single source electrode, since this extracellular electrode records from multiple neighbouring cells, the memristor is in fact tuned to detect the patterns of strong and synchronous network bursting activity. When there is no memristor-modulated communication between the source and target wells, the network activity of the six cultures are independent of each other (Figure 2B, left). When the memristor is inserted, the NBs of the source are detected due to the sudden increase of activity in the source electrode. This activity pattern automatically sets the memristor state to ON, closing the modulatory bridge between source and target wells via electrical stimulation. In turn, the electrical stimulus applied in the stimulation electrodes modulates neuronal activity in the target wells, thus establishing a communication pathway between previously isolated neuronal populations (Figure 2B, right). The short-term memory of the memristor operates at the temporal scales of the NB dynamics and guarantees that once the NB finishes in the source well, the memristor transitions back to OFF, avoiding the propagation of isolated neuronal spikes.

**Figure 2.**
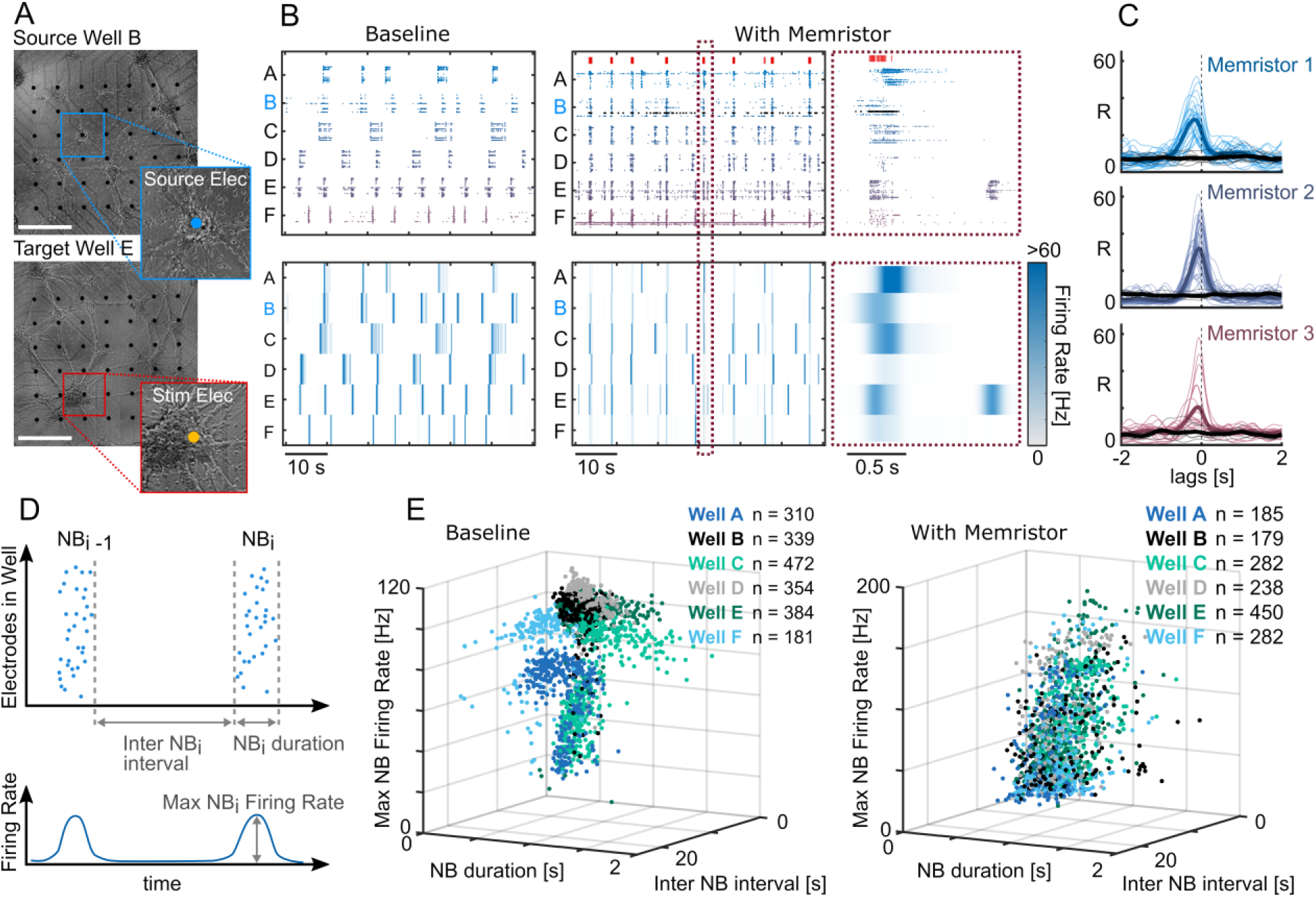
Real-time coupling and modulation of neuronal networks using a memristor. **A** Source and target neuronal networks cultured in different wells (scale bar = 500 μm). **B** Raster plot (top) and average firing rate per electrode on the distinct wells (bottom); the left and right panels correspond, respectively, to the system without and with the memristor-mediated connection. High frequency spikes in the source electrode (black dots, top right) correlate with network bursts in the source well (well B, in this representative example). **C** The synchronization of high-frequency activity between source and targets is evidenced by the cross-correlation (CC) curves between the average firing rate per electrode on the wells (B, bottom). There is a CC curve for each source-target pair per trial, for the three memristors tested (Memristor #1: 6 trials, 3.7±1.7 mins each; Memristor #2: 8 trials, 6.8±4.2 mins each; Memristor #3: 6 trials, 11.9±4.5 mins each). Coloured lines represent the CC curves for the trials with memristor, grey curves are for trials without memristor. Thicker lines represent the average. **D** Diagram describing the three parameters used for the network activity fingerprints: network burst (NB) duration, interval and maximum firing rate. **E** Memristive coupling modulates the dynamics of the network activity in the target wells, namely its NBs. Each dot represents a NB in a given well. Before coupling (left) the NBs of each well had consistent separable features (duration, interval, and maximum firing rate). After coupling, the network activity fingerprints between the source (well B) and target wells are indistinguishable (right) showing that the target wells’ neuronal populations are effectively responding to the source population activity.

Three different memristors were tested in separate trials, each monitoring a different source well. Qualitatively, the three memristors maintained a strong coupling between the high frequency activity patterns in the source and target wells when compared to the baseline (without memristor). To quantify this dynamic coupling, we computed the cross correlation (CC) between the mean firing rate of each source-target well pair, for each trial (**Figure 2C**). Despite the inherent variability of the memristor cycles, the CC curves showed a significant correlation between the firing rates of source and target wells, with a delay in the millisecond range. The memristive interface was thus capable of maintaining a robust low latency coupling of specific activity patterns between independent neuronal populations.

Besides quantifying the temporal coupling of the different networks, we also evaluated the impact of the memristive control on the modulation of activity fingerprints of the target networks. Specifically, we focused on the changes of properties such as duration and intervals between NBs, as well as maximum firing rates inside each NB (**Figure 2D**). In the system without the memristor, each well had each own, and independent, network activity fingerprint. In the memristor-based system, the network activity profile in all target wells followed, within physiological bounds, the command activity of the source well (**Figure 2E**). This shows that the dynamic modulation provided by the memristor aligned the activations of the target populations with the NBs of the source population. In the context of the detection-suppression system for epilepsy, the target neuronal population (or brain region) would be chosen as to have an inhibitory (or interfering) effect over the source population.

## 3 Conclusion

Neuromorphic devices are a promising alternative to overcome the balance between size, power and computational requirements, critical for innovative implantable neuroprosthesis. Memristors are neuromorphic devices that exhibit neuronal-like dynamics and can be tuned to operate in equivalent temporal scales. Here, we used memristors to interface with biological neuronal populations and selectively modulate their activity in a long-term autonomous setting. Our *in vitro* memristor-based system performs the three core tasks required for an efficient feedback neurostimulator - monitor, compute and actuate - and it does so with a micro-sized, low-power, neuromorphic computing element. The robust low latency detection and modulation of network activity patterns presented in this proof-of-concept would be highly desirable for an implantable neuronal stimulator in many clinically relevant contexts. Here, the computations were performed by a single memristor, allowing only for the detection of simple neuronal patterns. Future studies should focus on the integration of memristive arrays with biological neurons to perform real-time detection of more refined patterns of activity. Also, here the communication between neurons and memristors was mediated by an interface board and a conditioning circuit. Future interventions should focus on stablishing a direct/passive connection between neurons and memristors.

## 4 Experimental Section

### Memristive devices

The memristive devices used were acquired from *KNOWM Inc*. in the form of chips of 8 memristors and are composed of stacks of W/C+Ge_2_Se_2_/SnSe/Ge_2_Se_3_ Mix/Ag/Ge_2_Se_2_ Adhesion/W.^24^ They are analog devices with very low switching energy and fast switching response. The inherent threshold voltage of these devices varies in the range of approximately 0.25 - 0.45 V. To properly choose the values of the components in the electrical circuit (Figure S1), we performed a preceding characterization of the devices DC and pulse stimulation as can be seen in Figure 1C and Figure 1D.

### Cell culture

All experiments were performed in accordance with the European legislation for the use of animals for scientific purposes and protocols approved by the ethical committee of i3S. Embryonic (E18) rat hippocampal neurons were seeded and cultured at a density of 5 × 10 5 cells/well on a 6-well round chamber MEA with macrolon ring 10 mm high (256-6well MEA200/30iR-ITO-rcr) (Multichannel System MCS, Germany). The 6-well MEA electrode has an array of 7×6 TiN electrodes in each well, with a total of 252 recording electrodes. Half-medium changes of Neurobasal TM Plus (Thermo Fisher Scientific) were performed every 2-3 days.

### Electrophysiological recordings

The experiments were performed at 11 and 13 DIV using the MEA2100-256 system (Multichannel System MCS, Germany) at a sampling rate of 10 kHz. The electrophysiological signals were high-pass filtered at 200 Hz. The recordings were acquired using the Experimenter software from MCS. Cell culture conditions (37^a^C and 5% CO2) were maintained with a stage-top incubator (ibidi GmbH, Germany) adapted to the headstage of the MEA2100-256 system.

### Real-time spike detection

The detection of spikes in the source well was performed in real-time (latency of less than 20 μs) using the Digital Signal Processor (DSP) included in the interface board of the MEA2100-256 system. The spike detection was performed using a threshold crossing method on the already filtered signal. The negative threshold had 5 standard deviations (sometimes manually tuned at the beginning of the trials). The DSP was configured using the Experimenter software before starting the trials, being independent of the PC from there on. The interface board sent a TTL pulse to the memristor whenever the potential at the recording electrodes exceeds a pre-stablished threshold. The duration of the TTL was adjusted at the beginning of the trial to either 1, 5, 10 or 20 ms. Once configured, the entire system was fully independent of the PC.

### Mediating electrical circuit

The 3.3V TTL pulses arriving from the interface board were attenuated to fit the memristor operation range (millivolt range) using a dedicated circuit (see Supplementary Information). The circuit was tuned to ensure that only a high frequency of incoming pulses would set the memristor from OFF to ON in the required temporal scales. Also, the baseline level was set to a small negative value that ensured the transition from ON to OFF after a significant period without neuronal spikes. The signal processed by the memristors was converted back to the operating range of the Digital Input of the interface board.

### Electrical Stimulation

A stimulation electrode was selected for each target well. The electrical stimulus was a negative monophasic voltage pulse of 500mV with 200 μs. The electrical stimulation was triggered for all the stimulation electrodes every time the interface board received a high amplitude TTL pulse (above 2V) coming from the memristor. This way, when the memristor was ON, each spike detection in the source well triggered an electrical stimulation to each target well.

### Experimental Protocol

The experiments began with a baseline recording of 10-30 min, without memristor intervention. An electrode that had both bursting activity and isolated spikes was chosen as source electrode and its well, therefore, as source well. To select effective stimulation electrodes, the four most active electrodes of each well were individually stimulated with 20 electrical pulses with an interval of 10 secs. For each well, we chose the electrode that triggered a response in the largest number of neighbouring electrodes. The memristor was then inserted in the conditioning circuit connected to the interface board. Before starting the memristor experiment, the duration of the TTL pulses was adjusted to either 1, 5, 10 or 20 ms to assure proper short-term plasticity in the desired temporal scales. Each trial had a maximum duration of 15 min, but could be stopped earlier if the memristor was stuck in ON or OFF state for several minutes. The entirety of the recordings were considered in the analysis, including the periods where the memristor was apparently stuck in a given state.

### Signal Processing

The neuronal signals were filtered with a 200 Hz high-pass and spike detection was performed for all electrodes (not in real-time) using a positive and negative 6 standard deviations thresholds with 3 ms dead time. The mean firing rate of each well was obtained by convolution of the spike trains of the electrodes with a Gaussian kernel of 0.05 s sigma and averaging across electrodes. The cross-cross correlation curves were calculated between the mean firing rate profiles of each source-target well pairs. The network bursts (NB) consisted of periods when several electrodes bursted simultaneously. Events that included less than 5 bursting electrodes were not considered. To identify the bursting periods of each individual electrode associated with the NB, we considered the groups of at least 5 spikes with an interspike interval of less than 100 ms. The interspike interval considered for the burst detection in individual electrodes in larger than what is typically considered for bursting neurons (usually around 5 ms) because we wanted each electrode burst to encompass the full contribution of that electrode to the NB (instead of multiple bursts in the same electrode for the same NB).

## Acknowledgments

This work was financially supported by Project PTDC/EMD-EMD/31540/2017. The authors thank P. Cruz for all the help with the electrical circuit and its components.

## Supporting Information

**Figure S1.**
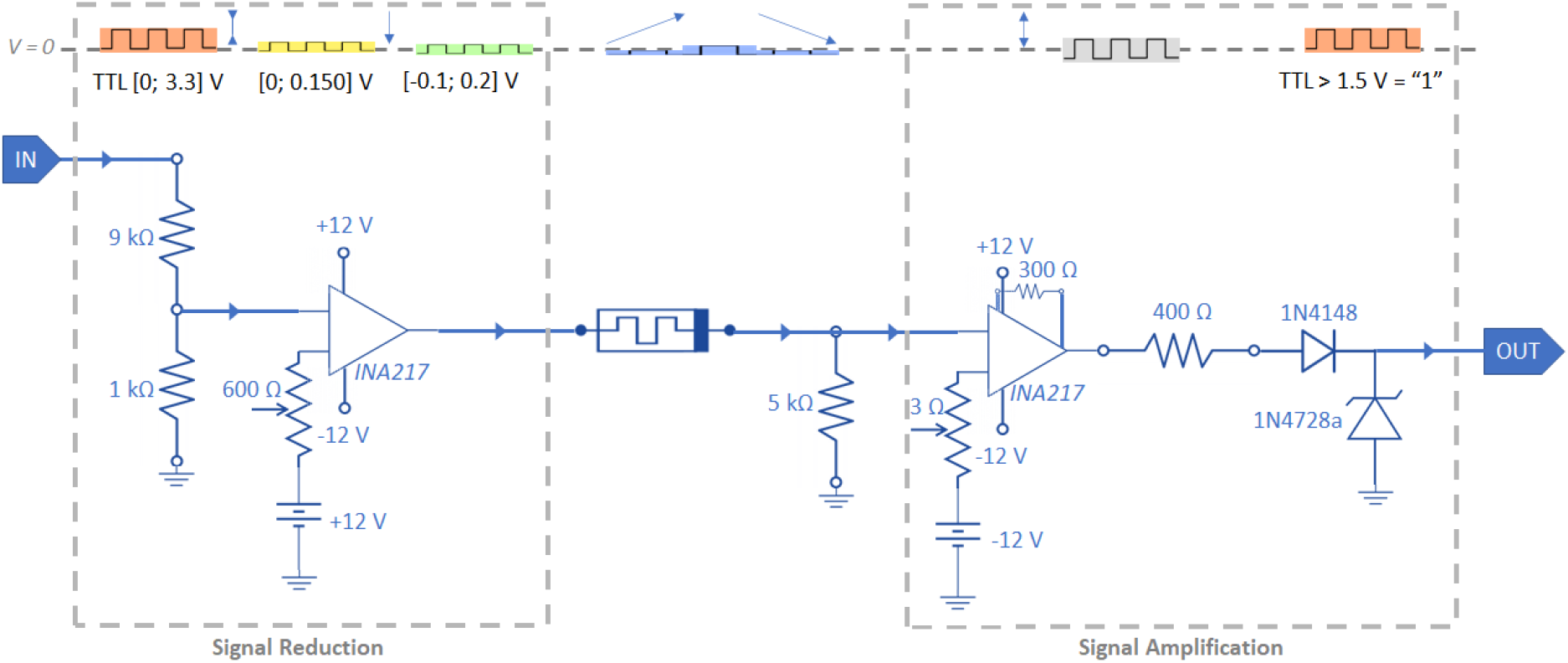
Detailed electric circuit used to process the signal between the digital input and the digital output of the MEA2100 interface board from MultiChannel Systems, showing the schematics of the signal across the circuit. The first block is responsible for the transformation of TTL signal to the lower operation voltage range of the memristor, using a voltage divider and a low-noise instrumentation amplifier to shift the pulse to have both a positive (for “set”) and a negative (for “reset”) component. The last block amplifies the signal at the memristor output, for it to be read as a “0” when the memristor is OFF and “1” when the memristor is ON. The diodes are only present as a precaution for protection of the data acquisition system.

